# The pre-Bötzinger complex is necessary for the expression of post-inspiration

**DOI:** 10.1101/2022.03.28.486134

**Authors:** Rishi R. Dhingra, Werner I. Furuya, Mathias Dutschmann

## Abstract

The mammalian three-phase respiratory motor pattern of inspiration, post-inspiration and expiration is expressed in spinal and cranial motor nerve activities. This pattern is generated by a hierarchical brainstem-wide pre-motor network. However, the local rhythmogenic circuit of the pre-Bötzinger complex (pre-BötC) was established as the ‘noeud vitale’ necessary and sufficient to initiate inspiration. In present study, we tested the effect of unilateral and bilateral inactivation of the pre-BötC (microinjection of the GABA_A_ receptor agonist isoguvacine 10 mM, 50 nl) on respiratory motor activities in an *in situ* perfused brainstem preparation of rat. As expected, bilateral inactivation of the pre-BötC triggered cessation of phrenic (PNA), hypoglossal (HNA) and vagal (VNA) nerve activities for 15-20 min. Partial recovery from bilateral isoguvacine injection was characterized by erratic activity in all recorded motor nerves reminiscent to patterns observed after disturbed excitatory-inhibitory balance within the local pre-BötC circuit. Surprisingly, ipsilateral isoguvacine injections into the pre-BötC triggered transient (6-8 min) cessation of inspiratory and post-inspiratory VNA (p<0.001) and suppressed inspiratory HNA by -70 ± 15% (p<0.01), while inspiratory PNA burst frequency increased by 46 ± 30% (p<0.01). Taken together, these observations confirm the role of the pre-BötC as the ‘noeud vitale’ of the mammalian respiratory network *in situ* and highlight a significant role for the pre-BötC in the expression of vagal post-inspiratory and inspiratory activity.

## Introduction

A key discovery to understand the neural control of breathing was the identification of the pre-Bötzinger complex (pre-BötC), a localized nucleus in the rostral medulla oblongata that generates the inspiratory rhythm *in vitro* (Smith *et al*., 1991). Importantly, elegant lesioning of the pre-BötC in the intact respiratory network established its role as the “noeud vitale” that is sufficient and essential for the generation of the vital inspiratory rhythm (Gray *et al*., 2001; McKay *et al*., 2005; Tan *et al*., 2008). Therefore, many studies have investigated the mechanisms underlying the intrinsic bursting of pre-BötC neurons experimentally using *in vitro* slice preparations and computationally (Thoby-Brisson & Ramirez, 2001; Del Negro *et al*., 2002,2005; Rybak *et al*., 2003; Peña *et al*., 2004; Shao *et al*., 2006; Rubin *et al*., 2009; Kam et al., 2013a,b; Rubin & Smith, 2019; Ashhad & Feldman, 2020).

However, under intact network conditions, the respiratory motor pattern consists of three phases of activity—inspiration, post-inspiration and late-expiration—that are observed on spinal and cranial nerves (Dutschmann & Paton, 2002). The generation of this three-phase motor pattern is thought to depend on mutual inhibition between medullary neuronal populations that share these bursting patterns (Richter 1982; Richter & Spyer, 2001; Richter & Smith, 2014). Reconciling these observations, computational models of respiratory pattern generation now contain intrinsically bursting pre-inspiratory neurons of the pre-BötC (Rybak *et al*., 2004; Smith et al., 2007, 2013; Molkov *et al*., 2016; Ausborn *et al*., 2018), a feature whose necessity for respiratory pattern generation has also been contested (Rubin & Smith, 2019).

Recently, experimental data from our laboratory and others have shown that the pre-BötC not only plays a role in the generation of the inspiratory rhythm, but also engages with an anatomically distributed network necessary for the expression of the three-phase respiratory motor pattern. Local disruption of excitatory-inhibitory balance in the pre-BötC under intact network conditions in the *in situ* perfused brainstem preparation, while permitting the generation of the respiratory rhythm, pathologically disrupts the three-phase respiratory motor pattern (Dhingra *et al*., 2019a,b). Further, recent measurements of respiratory local field potentials in the pre-BötC *in situ* demonstrated the robust expression of LFPs at the transition from expiration to inspiration and from inspiration to post-inspiration (Dhingra *et al*., 2020). Finally, optogenetic activation of a subset of pre-BötC neurons *in vivo* showed phase-dependent effects such that activation of pre-BötC neurons during inspiration increases respiratory frequency, whereas activation during post-inspiration depresses respiratory frequency (Hülsmann *et al*., 2021). Taken together, these data suggest a novel role for the pre-BötC in determining the timing of the inspiratory/post-inspiratory phase transition.

Therefore, in the present study, we investigated the hypothesis that the pre-BötC is not only necessary for the generation of the respiratory rhythm, but also serves as a critical component of a distributed post-inspiratory circuit, whose integrity is necessary for the expression of the three-phase respiratory motor pattern. We tested our hypothesis by examining the effects of local pharmacologic inactivation of the pre-BötC in the *in situ* perfused brainstem preparation. As expected from previous work, bilateral inactivation of the pre-BötC transiently ceased all respiratory motor activities. However, unilateral inactivation of the pre-BötC transiently suppressed vagal post-inspiratory and inspiratory activities with only a minor effect on diaphragmatic motor output. Together, these data provide further evidence that the pre-BötC is necessary for the expression of post-inspiratory motor activity.

## Materials and Methods

Experimental protocols were approved by and conducted with strict adherence to the guidelines established by the Animal Ethics Committee of the Florey Department of Neuroscience & Mental Health, University of Melbourne, Melbourne, Australia.

### 2.1 In situ arterially-perfused brainstem preparation

Experiments were performed in juvenile (17-26 days post-natal) Sprague-Dawley rats of either sex using the *in situ* arterially-perfused working heart brainstem preparation without the heart, as described previously (Paton, 1996; Paton *et al*., 2022). Briefly, rats were anesthetized by inhalation of isoflurane (2-5%) until they reached a surgical plane of anesthesia. Next, rats were transected sub-diaphragmatically and immediately transferred to an ice-cold bath of artificial cerebrospinal fluid (aCSF; in mM: [NaCl] 125, [KCl] 3, [KH_2_PO_4_] 1.25, [MgSO_4_] 1.25, [NaHCO_3_] 24, [CaCl_2_] 2.5 and D-glucose 10) for decerebration. Next, the heart and lungs were removed. The phrenic nerve was isolated for recording, and the descending aorta was isolated for cannulation. Next, the cerebellum was removed. Finally, the vagus and hypoglossal nerves were isolated for recording. The preparation was then transferred to a recording chamber. The aorta was quickly cannulated with a double-lumen catheter. The preparation was then re-perfused with carbogenated (95%/5% pO_2_/pCO_2_), heated (31°C) aCSF (200 mL) using a peristaltic pump (Watson-Marlow).

All recorded nerves were mounted on suction electrodes to measure respiratory motor output. Nerve potentials were amplified (10,000×, Warner Instruments, DP-311), band-pass filtered (0.1–10 kHz), digitized (AD Instruments, PowerLab 16/35), and stored on a computer using LabChart software (AD Instruments version 7.0). Within minutes, apneustic respiratory contractions resumed.

A single bolus of NaCN (0.1 mL, 0.1% (w/v) provided excitatory chemosensory drive to convert the apneustic pattern to a eupneic three-phase respiratory motor pattern. The perfusion flow rate was adjusted to fine tune the preparation to generate a stable stationary rhythm. Finally, a single bolus of vecuronium bromide (0.3 mL, 0.1 mg/mL w/v vecuronium bromide: saline) was delivered to the perfusate to paralyse the preparation to avoid movement artifacts.

### 2.2 Pre-BötC inhibition with isoguvacine

We locally micro-injected the GABA_A_R agonist isoguvacine (50nL, 10mM in aCSF) in the pre-BötC and measured the effect of pre-BötC inhibition on the three-phase respiratory motor pattern as expressed across phrenic, hypoglossal and vagal nerves. For these experiments, we used a triple-barreled pipette that also included a barrel filled with glutamate (10mM in aCSF) and another filled with pontamine sky blue (0.1% w/v in aCSF) to enable functional mapping of the injection site and *post-hoc* verification of the injection sites, respectively. Injection volumes were monitored using a calibrated reticule. Glutamate microinjection (50 nL) into the pre-BötC evoked a brief tachypnea. Effective pre-BötC coordinates were: 1.4-1.5 mm rostral to *calamus scriptorius*, 1.8-1.9 mm lateral to the midline and 2.0 mm below the brainstem surface. Once the pre-BötC was functionally identified, we microinjected isoguvacine (50nL) unilaterally at those sites. We then recorded the respiratory motor pattern after we microinjected isoguvacine unilaterally for 6-8 minutes, before microinjecting isoguvacine at the same coordinates in the contralateral hemisphere. The effect of bilateral pre-BötC inhibition was monitored for at least 20 minutes. To determine the noise levels, the perfusion was ceased at the end of experiments and the recording was sustained for additional 10 min following the complete silencing of the nerves.

At the conclusion of the experiment, the brainstem was removed and post-fixed in 4% paraformaldehyde for 3-5 days, and subsequently, cryo-protected in 0.1M PBS containing 20% sucrose for 2 days. Brainstems were cryo-sectioned (Leica CM1900, 40 µm thickness), mounted and counterstained with 1% Neutral Red (Sigma-Aldrich Cat No. 861251) to verify that isoguvacine injection sites were within the anatomical location of the pre-BötC.

### 2.3 Pre-BötC inhibition with isoguvacine – Data Analysis

Analysis of respiratory motor activity were performed with Spike2 software version 7.17 (Cambridge Electronic Design, Cambridge, UK). All analyses were performed in 6 distinct 1-min intervals: baseline, 1-2 min after unilateral injection of isoguvacine in the pre-BötC and at 2, 5, 10 and 20 min following the bilateral injection. Phrenic, vagal and hypoglossal nerve activities (PNA, VNA & HNA, respectively) were rectified and integrated (50 ms time constant).

The respiratory frequency (*F*_R_) was determined by the time interval between the onset of inspiration of consecutive bursts as reflected in the integrated PNA. To assess the effect of pre-BötC inhibition on respiratory motor nerve amplitude, we measured the area under the curve (AUC) of the integrated waveform of PNA, VNA and HNA, separately. The AUC was calculated as the total area of the bursts observed within the 1-min interval analyzed, after noise subtraction. The noise threshold was determined from the period following the perfusion cessation at the end of experiment. To test the significance of these effects, we used a one-way repeated measures ANOVA followed by Student-Newman-Keuls *post-hoc* test to identify specific differences.

## Results

### The pre-Bötzinger complex—the ‘noeud vitale’ of the intact respiratory network

Consistent with previous studies (Gray *et al*., 2001; McKay *et al*., 2005; Tan *et al*., *2008*), bilateral pre-BötC inhibition (Fig. 1) terminated respiratory activity. Within 2 minutes after bilateral pre-BötC inhibition, we observed the complete cessation of rhythmic bursting (15.4 ± 3.3 bursts/min to 0.0 ± 0.0 bursts/min; *p<0.001*, Fig. 1A1-2 & B) in all recorded respiratory cranial (HNA & VNA) and spinal nerves (PNA), confirming the role of the pre-BötC as the ‘noeud vitale’ of the intact respiratory network *in situ*. Nevertheless, bursting activities in respiratory motor outputs often re-emerged within 10-15 min after bilateral isoguvacine injections (Fig. 1A3 & B).

**Figure 1:**
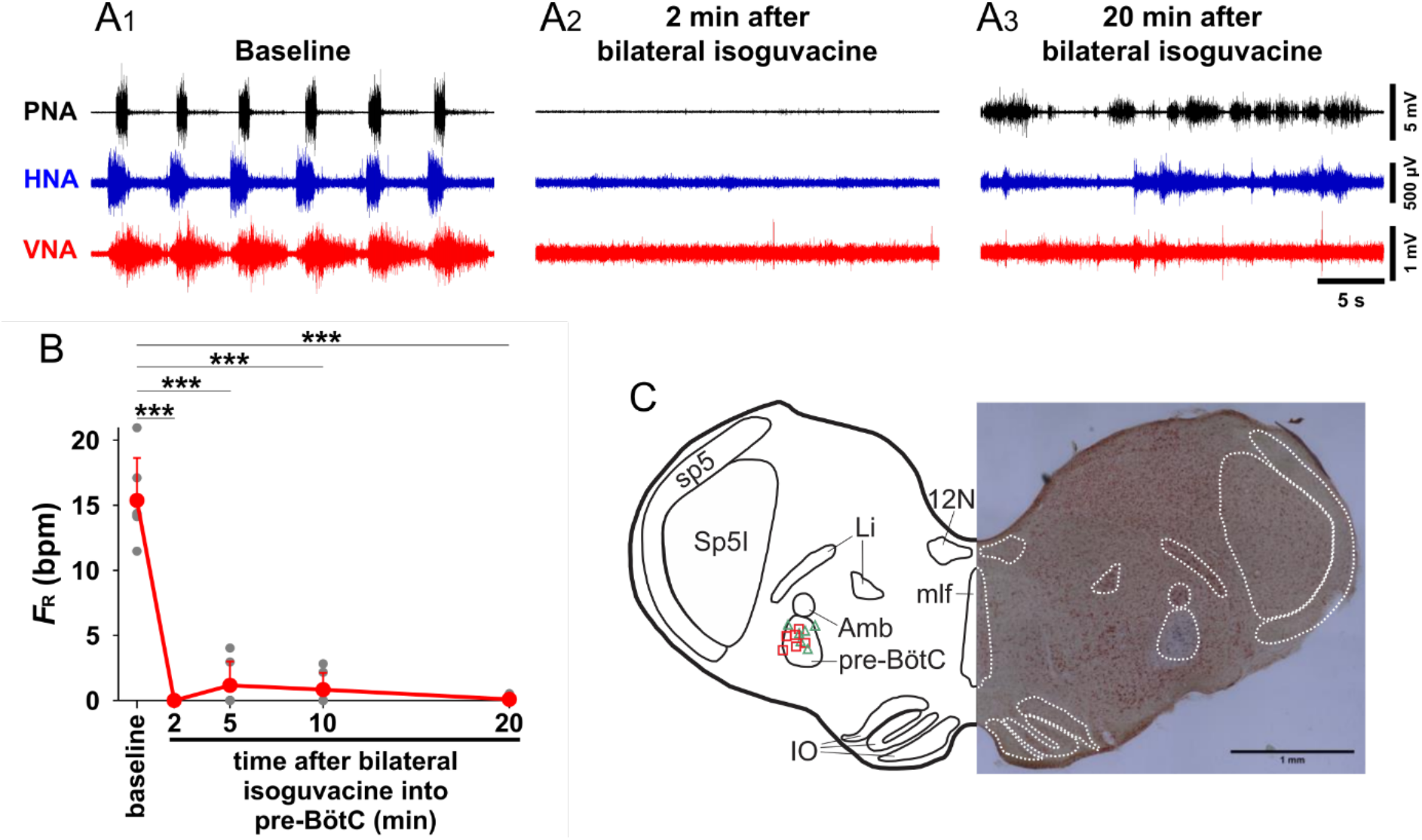
The pre-Bötzinger Complex (pre-BötC) is the ‘noeud vitale’ of the respiratory network in the in situ perfused preparation. **A** Representative recording of phrenic (PNA), hypoglossal (HNA) and vagal (VNA) nerve activity at baseline (A1), 2 min (A2) and 20 min (A3) after bilateral injection of isoguvacine into the pre-BötC. The bilateral inhibition of the pre-BötC abolished the eupneic respiratory pattern **B** The eupneic phrenic burst frequency (FR) remained suppressed for at least 20 min following the isoguvacine injections into the pre-BötC. *** p< 0.001, n=6 **C** Schematic drawing and photomicrograph of coronal section of the brainstem indicating the microinjection sites of isoguvacine in the pre-BötC. Green triangles denote left hemisphere microinjections, whereas red squares denote right hemisphere microinjections. 12N, hypoglossal nucleus; Amb, nucleus ambiguus; IO, inferior olive; Li, linear nucleus of the medulla; mlf, medial longitudinal fasciculus; pre-BötC, pre-Bötzinger complex; py, pyramidal tract; sp5, spinal trigeminal tract; Sp5I, interpolar spinal trigeminal nucleus.

However, these bursts were of a low-amplitude and did not generate a three-phase respiratory motor pattern. Thus, they were not scored as eupneic breaths in our analysis of pre-BötC inhibition effects on respiratory frequency. These pathologic motor patterns were reminiscent with those observed after perturbing excitation-inhibition balance by increasing excitability via local application of the GABA_A_R antagonist bicuculline in the pre-BötC (Dhingra *et al*., 2019a, 2019b).

### The pre-Bötzinger Complex is an integral part of the cranial nerve motor system

Next, we focused on the effect of unilateral pre-BötC inhibition on spinal and cranial nerve discharge patterns (Fig. 2). This analysis revealed that unilateral inhibition of pre-BötC evoked larger effects on HNA and VNA compared to the PNA (Fig. 2B). Within 30 seconds, unilateral pre-BötC inactivation abolished VNA (Fig. 2B) and significantly reduced the amplitude of HNA (0.4 ± 0.2 vs. baseline 1.5 ± 0.4 V ·s,*p<0.01*, Fig. 2B), but only partially reduced the amplitude of PNA (3.6 ± 2.7 vs. baseline 4.1 ± 2.3 V·s, *p>0.05*) Fig. 2A1-2).

**Figure 2:**
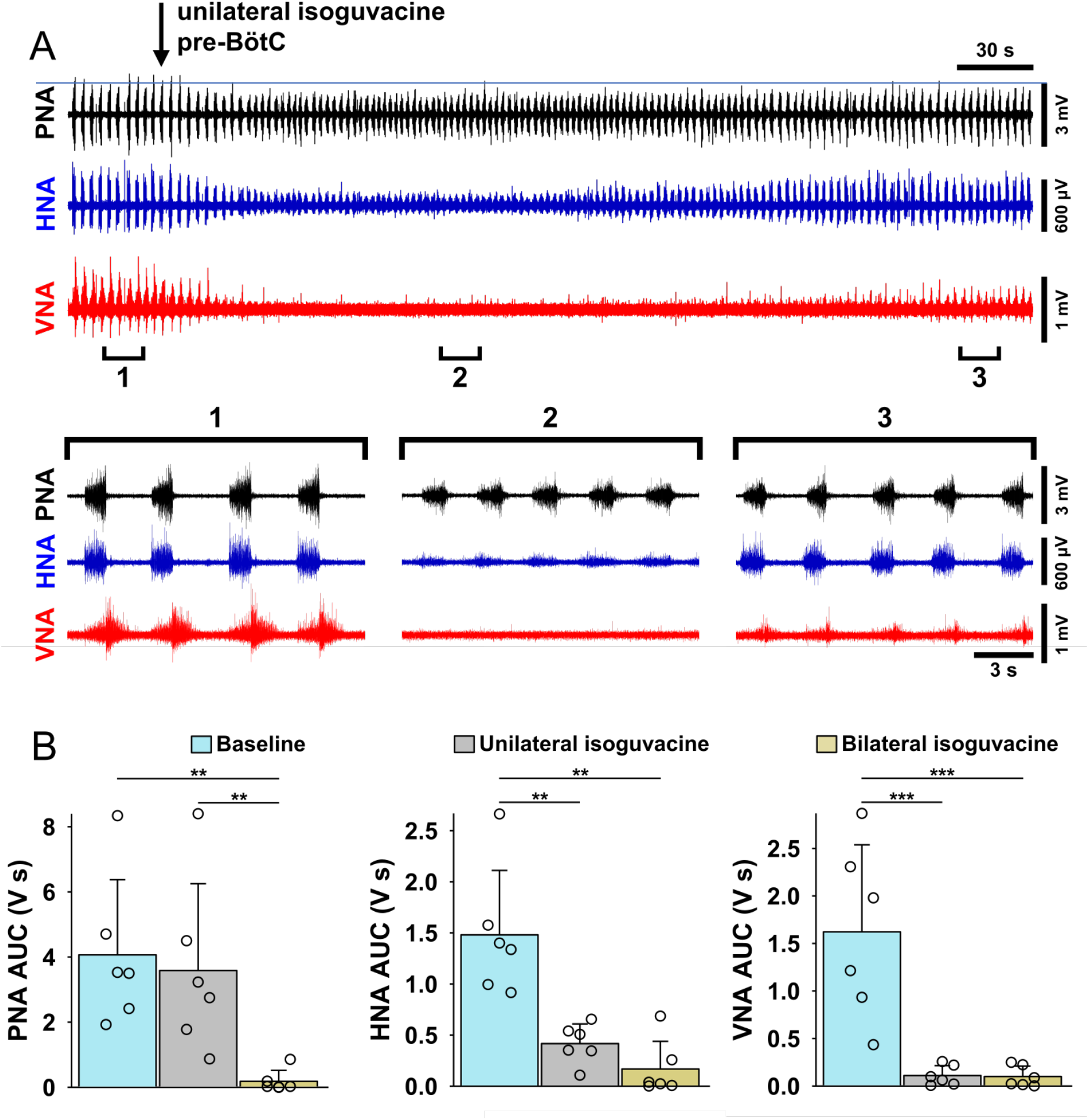
The cranial respiratory motor pattern depends on the integrity of the pre-BötC. **A** Representative recordings of phrenic (PNA), hypoglossal (HNA) and vagal (VNA) nerve activity at baseline (1), 2 min (2) and 5 min (3) after unilateral injection of isoguvacine into the pre-BötC. The unilateral inhibition of the pre-BötC induced a substantial decrease in the HNA amplitude and completely abolished the VNA. The eupneic respiratory motor pattern partially recovered 5 min after the injection (3) **B** The area under the curve (AUC) obtained from the integrated traces demonstrates that unilateral inhibition of the pre-BötC was sufficient to suppress HNA and VNA, but not PNA. The eupneic respiratory nerve amplitudes were abolished following bilateral inhibition of the pre-BötC *** *p* < 0.001, ** *p* < 0.01, n=6.

In addition, PNA burst frequency (respiratory rate) increased (22.8 ± 7.4 vs. baseline 15.4 ± 3.3 bursts/min, *p<0.01*) after unilateral inhibition of the pre-BötC (Fig. 1B). Within 5-8 minutes, VNA and HNA motor patterns recovered to control (Fig. 2A3). However, VNA amplitude remained reduced compared to control (0.2 ± 0.1 vs. baseline 1.6 ± 0.9 V · s, *p<0.001*). Taken together, the effects of unilateral pre-BötC inhibition suggests that the pre-BötC serves as an integral component of the highly-distributed post-inspiratory circuit.

## Discussion

In the present study, we showed that bilateral inhibition of the pre-BötC in the intact respiratory network of the *in situ* perfused preparation suppresses all respiratory motor output on cranial and spinal nerves consistent with the view of the pre-BötC as the ‘noued vitale’ of the respiratory network (Smith *et al*., 1991; Gray *et al*., 2001; McKay *et al*., 2005; Tan *et al*., 2008). Surprisingly, unilateral inhibition of the ipsilateral pre-BötC *in situ* was sufficient to abolish inspiratory and post-inspiratory activity of the vagus nerve and to depress hypoglossal nerve output. The latter observation highlights a novel role of the pre-BötC in the regulation of post-inspiration and hence upper airway patency during resting breathing.

The present study shows for the first time that unilateral pre-BötC inhibition silences both inspiratory and post-inspiratory output in the ipsilateral vagal nerve, which mediates the laryngeal valving of respiratory airflow (see Paton & Dutschmann, 2002). This observation is functionally consistent with recent experiments in which optogenetic activation of pre-BötC neurons during expiration prolongs expiration (Hülsmann *et al*., 2021). The observation that unilateral inhibition of the pre-BötC causes suppression of inspiratory hypoglossal motor output is in line with previous reports that pre-BötC activity specifically drives the inspiratory activity of the hypoglossal nerve (Smith *et al*., 1991, Del Negro *et al*., 2002, 2005). The suppression hypoglossal nerve activity causes diminished genioglossus protrusion and retraction during inspiration (Fuller *et al*., 1999; Gestreau *et al*., 2005, Lee *et al*., 2012). Together, these findings suggest that the pre-BötC contains a critical pool of pre-motor neurons that project to hypoglossal and vagal motor neurons (Guo *et al*., 2020; Menuet *et al*., 2020), which have an essential function in the dynamic regulation of upper airway patency. Thus, the present study further extends the role of the pre-BötC from its established role as the inspiratory rhythm generator (Feldman & Del Negro 2006; Feldman *et al*., 2008; Del Negro *et al*., 2018) to the dynamic regulation of upper airway patency and is also consistent with its recently identified role in the mediation of cough (Shen et al., 2022) that specifically requires post-inspiratory laryngeal constriction to generate high velocity expiratory airflow (see Dutschmann et al., 2014).

The notion that the pre-BötC circuit is important for the regulation of post-inspiration adds to a growing list of brainstem areas involved in the control of this phase of the respiratory cycle. In particular, the contribution of the pontine Kölliker-Fuse nucleus to post-inspiration, and to the gating of cranial nerve activities, was previously highlighted by our laboratory (Dutschmann & Herbert, 2006; Bautista & Dutschmann, 2014; Dutschmann *et al*., 2021). With respect to the post-inspiratory phase, others have pointed to the recently coined post-inspiratory complex (PiCo, Anderson *et al*., 2016) or the Bötzinger complex (Burke *et al*., 2008) as key medullary areas for the generation of post-inspiratory activity. However, the assertion that any one brainstem nucleus solely determines post-inspiratory activity is flawed. *A priori* transected brainstems that lack the pons, but maintain medullary connectivity including that between the PiCo, the BötC and pre-BötC fail to generate a three-phase respiratory motor pattern (Jones & Dutschmann, 2016) suggesting that connectivity within the distributed ponto-medullary respiratory network determines the vagal inspiratory/post-inspiratory motor pattern. Similarly, in the context of the present report, it would be inaccurate to suggest that the pre-BötC is the sole determinant of post-inspiratory activity since recent optogenetic inactivation of pre-BötC neurons evoked enhanced post-inspiratory VNA (Menuet *et al*., 2020). Instead, post-inspiratory activity appears to be determined by a distributed network. Accordingly, local disruption of activity within any of these compartments disrupts the three-phase respiratory motor pattern despite sparing the remaining connectivity of the respiratory network (Dhingra *et al*., 2019a,b). We recently confirmed this view of a distributed circuit mediating the transition to post-inspiration using a newly developed approach to map respiratory LFPs at hundreds of sites across the brainstem in single *in situ* preparations using silicon multi-electrode arrays (Dhingra *et al*., 2020). These data provided compelling support that widely-distributed network areas in the pons, and in dorsal and ventral medullary areas act cooperatively to engage the transition into the post-inspiratory phase.

The recruitment of distributed post-inspiratory neurons during orofacial behaviors such as swallowing, vocalization or sniffing, which all require specific modulations of airway patency, is further supported by recent trans-synaptic tracing studies that identified a brainstem-wide pre-motor network that overlaps with key nodes of the primary respiratory pattern generation network, including the Kölliker-Fuse nucleus, BötC, pre-BötC and intermediate reticular nucleus (Kurnikova *et al*., 2018; Takatoh *et al*., 2021). Further, both the pre-BötC and the Kölliker-Fuse nucleus are specific targets of descending cortical projections (Yang *et al*., 2020; Trevizan-Baú *et al*., 2021) emphasizing that key nodes for the coordination of volitional orofacial behaviors with on-going breathing activities are likely to be anatomically distributed. Thus, the synaptic interplay of an extended and distributed brainstem circuit is required for the generation of cranial nerve motor patterns during eupnea as well as during the coordination of breathing with vital orofacial behaviors.

## Conclusion

We conclude that in addition to being the ‘noeud vitale’ of the respiratory network *in situ*, the pre-BötC is a necessary component of the distributed network that underlies the expression of post-inspiratory activity in the vagal nerve. The latter is critical for the control of upper airway patency during resting breathing and for the adaptation of the breathing pattern during respiratory reflexes and orofacial behaviour.

## Acknowledgements

We acknowledge that this work was conducted on the traditional land of the Wurundjeri people of the Kulin nation. We pay our respect to their elders past, present and emerging. This work was supported by research grants from the National Health and Medical Research Council of Australia (APP1129376); and Australian Research Council (DP170104861).

## References

Anderson TM, Garcia AJ 3rd, Baertsch NA, Pollak J, Bloom JC, Wei AD, Rai KG & Ramirez JM (2016). A novel excitatory network for the control of breathing. Nature 536(7614), 76–80. doi: 10.1038/nature18944.

Ashhad S & Feldman JL (2020). Emergent Elements of Inspiratory Rhythmogenesis: Network Synchronization and Synchrony Propagation. Neuron 106(3), 482–497. doi: 10.1016/j.neuron.2020.02.005. Epub 2020 Mar 3. PMID: 32130872.

Ausborn J, Koizumi H, Barnett WH, John TT, Zhang R, Molkov YI, Smith JC & Rybak IA (2018). Organization of the core respiratory network: Insights from optogenetic and modeling studies. PLoS Computational Biology. 14(4), e1006148. doi: 10.1371/journal.pcbi.1006148.

Bautista TG & Dutschmann M (2014). Inhibition of the pontine Kölliker-Fuse nucleus abolishes eupneic inspiratory hypoglossal motor discharge in rat. Neuroscience 267, 22–9. doi: 10.1016/j.neuroscience.2014.02.027.

Burke PG, Abbott SB, McMullan S, Goodchild AK & Pilowsky PM (2010). Somatostatin selectively ablates post-inspiratory activity after injection into the Bötzinger complex. Neuroscience. 167(2), 528–39. doi: 10.1016/j.neuroscience.2010.01.065.

Del Negro CA, Morgado-Valle C & Feldman JL (2002). Respiratory Rhythm: An Emergent Network Property? Neuron 34, 821–830.

Del Negro CA, Morgado-Valle C, Hayes JA, Mackay DD, Pace RW, Crowder EA & Feldman JL (2005). Sodium and Calcium Current-Mediated Pacemaker Neurons and Respiratory Rhythm Generation. J Neurosci 25, 446–453.

Del Negro CA, Funk GD & Feldman JL (2018). Breathing matters. Nature Reviews Neuroscience. 19(6), 351–367. doi: 10.1038/s41583-018-0003-6.

Dhingra RR, Dick TE, Furuya WI, Galán RF & Dutschmann M (2020). Volumetric mapping of the functional neuroanatomy of the respiratory network in the perfused brainstem preparation of rats. J Physiol 598, 2061–2079.

Dhingra RR, Furuya WI, Bautista TG, Dick TE, Galán RF & Dutschmann M (2019a). Increasing Local Excitability of Brainstem Respiratory Nuclei Reveals a Distributed Network Underlying Respiratory Motor Pattern Formation. Front Physiol; DOI: 10.3389/fphys.2019.00887.

Dhingra RR, Furuya WI, Galán RF & Dutschmann M (2019b). Excitation-inhibition balance regulates the patterning of spinal and cranial inspiratory motor outputs in rats in situ. Respir Physiol Neurobiol 266, 95–102.

Dutschmann M & Herbert H (2006). The Kölliker-Fuse nucleus gates the postinspiratory phase of the respiratory cycle to control inspiratory off-switch and upper airway resistance in rat. European Journal of Neuroscience 24, 1071–1084.

Dutschmann M, Jones SE, Subramanian HH, Stanic D, & Bautista TG (2014). The physiological significance of postinspiration in respiratory control. Progress in Brain Research 212, 113–30. doi: 10.1016/B978-0-444-63488-7.00007-0.

Dutschmann M, Bautista TG, Trevizan-Baú P, Dhingra RR, & Furuya WI (2021). The pontine Kölliker-Fuse nucleus gates facial, hypoglossal, and vagal upper airway related motor activity. Respiratory Physiology & Neurobiology 284, 103563. doi: 10.1016/j.resp.2020.103563.

Feldman JL & Del Negro CA (2006). Looking for inspiration: new perspectives on respiratory rhythm. Nature Reviews Neuroscience 7(3), 232–42. doi: 10.1038/nrn1871. PMID: 16495944; PMCID: PMC2819067.

Feldman JL, Del Negro CA & Gray PA (2013). Understanding the rhythm of breathing: so near, yet so far. Annual Reviews of Physiology. 75, 423–52. doi: 10.1146/annurev-physiol-040510-130049.

Fuller DD, Williams, JS, Janssen, PL, Fregosi, R.F., 1999. Effect of co-activation of tongue protrudor and retractor muscles on tongue movements and pharyngeal airflow mechanics in the rat. The Journal of Physiology 519, 601–613.

Gestreau C, Dutschmann M, Obled S & Bianchi AL (2005). Activation of XII motoneurons and premotor neurons during various oropharyngeal behaviors. Respiratory Physiology & Neurobiology 147(2-3), 159–76. doi: 10.1016/j.resp.2005.03.015. PMID: 15919245.

Guo H, Yuan XS, Zhou JC, Chen H, Li SQ, Qu WM & Huang ZL (2020). Whole-Brain Monosynaptic Inputs to Hypoglossal Motor Neurons in Mice. Neuroscience Bulletin 36(6), 585–597. doi: 10.1007/s12264-020-00468-9.

Gray PA, Janczewski WA, Mellen N, McCrimmon DR, Feldman JL (2001). Normal breathing requires preBötzinger complex neurokinin-1 receptor-expressing neurons. Nat Neurosci 4, 927–30. doi: 10.1038/nn0901-927.

Hülsmann S, Hagos L, Eulenburg V, Hirrlinger J (2021). Inspiratory Off-Switch Mediated by Optogenetic Activation of Inhibitory Neurons in the preBötzinger Complex In Vivo. Int J Mol Sci 22,2019. doi: 10.3390/ijms22042019.

Jones SE & Dutschmann M (2016). Testing the hypothesis of neurodegeneracy in respiratory network function with a priori transected arterially perfused brain stem preparation of rat. J Neurophysiol 115, 2593–2607.

Kam K, Worrell JW, Janczewski WA, Cui Y & Feldman JL (2013a). Distinct Inspiratory Rhythm and Pattern Generating Mechanisms in the preBötzinger Complex. J Neurosci 33, 9235–9245.

Kam K, Worrell JW, Ventalon C, Emiliani V & Feldman JL (2013b). Emergence of Population Bursts from Simultaneous Activation of Small Subsets of preBötzinger Complex Inspiratory Neurons. J Neurosci 33, 3332–3338.

Kurnikova A, Deschenes M & Kleinfeld D (2018). Functional brainstem circuits for control of nose motion. Journal of Neurophysiology 121(1),205–217. DOI: 10.1152/jn.00608.2018.

Lee KZ, Fuller DD & Hwang JC (2012). Pulmonary C-fiber activation attenuates respiratory-related tongue movements. Journal of Applied. Physiology (1985) 113, 1369–1376.

McKay LC, Janczewski WA, Feldman JL (2005). Sleep-disordered breathing after targeted ablation of preBötzinger complex neurons. Nat Neurosci 8, 1142–4. doi: 10.1038/nn1517.E

Menuet C, Connelly AA, Bassi JK, Melo MR, L. S, Kamar J, Kumar NN, McDougall SJ, McMullan S, Allen AM (2020). PreBötzinger complex neurons drive respiratory modulation of blood pressure and heart rate. Elife 9:e57288. doi: 10.7554/eLife.57288.

Molkov YI, Rubin JE, Rybak IA & Smith JC (2017). Computational models of the neural control of breathing. Wiley Interdisciplinary Reviews: System Biology and Medicine. 9(2),10.1002/wsbm.1371. doi: 10.1002/wsbm.1371.

Paton JFR (1996). A working heart-brainstem preparation of the mouse. Journal of Neuroscience Methods 65, 63–68.

Paton JFR, Machado, BH, Moraes DJA, Daniel B Zoccal DB, Abdala AP, Smith JC, Vagner Roberto Antunes VR, Murphy D, Dutschmann M, Dhingra RR, McAllen RM, Pickering AE, Wilson RJA, Day TA, Barioni NO, Allen AM, Menuet C, Donnelly J, Felippe ISA, & St John WM. Advancing respiratory-cardiovascular physiology with the working heart-brainstem preparation over 25 years. J Physiol, doi: 10.1113/JP281953

Paton JF & Dutschmann M (2002). Central control of upper airway resistance regulating respiratory airflow in mammals. Journal of Anatomy 201(4), 319–23. doi: 10.1046/j.1469-7580.2002.00101.x.

Peña F, Parkis MA, Tryba AK & Ramirez J-M (2004). Differential Contribution of Pacemaker Properties to the Generation of Respiratory Rhythms during Normoxia and Hypoxia. Neuron 43, 105–117.

Richter DW (1982). Generation and maintenance of the respiratory rhythm. Journal of Experimental Biology 100, 93–107. doi: 10.1242/jeb.100.1.93. PMID: 6757372.

Richter DW & Spyer KM (2001). Studying rhythmogenesis of breathing: comparison of in vivo and in vitro models. Trends in Neuroscience 24(8), 464–72. doi: 10.1016/s0166-2236(00)01867-1.

Richter DW & Smith JC (2014). Respiratory rhythm generation in vivo. Physiology (Bethesda). 29(1), 58–71. doi: 10.1152/physiol.00035.2013.

Rubin JE, Shevtsova NA, Ermentrout GB, Smith JC & Rybak IA (2009). Multiple Rhythmic States in a Model of the Respiratory Central Pattern Generator. Journal of Neurophysiology 101, 2146–2165.

Rubin JE & Smith JC (2019). Robustness of respiratory rhythm generation across dynamic regimes. PLOS Computational Biology 15, e1006860.

Rybak IA, Shevtsova NA, St-John WM, Paton JFR & Pierrefiche O (2003). Endogenous rhythm generation in the pre-Bötzinger complex and ionic currents: modelling and in vitro studies. European Journal of Neuroscience 18, 239–257.

Rybak IA, Shevtsova NA, Paton JFR, Dick TE, St.-John WM, Mörschel M & Dutschmann M (2004). Modeling the ponto-medullary respiratory network. Respiratory Physiology & Neurobiology 143, 307–319.

Shao J, Tsao T-H & Butera R (2006). Bursting Without Slow Kinetics: A Role for a Small World? Neural Computation 18, 2029–2035.

Shen TY, Poliacek I, Rose MJ, Musselwhite MN, Kotmanova Z, Martvon L, Pitts T, Davenport PW, Bolser DC(2022). The role of neuronal excitation and inhibition in the pre-Bötzinger complex on the cough reflex in the cat. Journal of Neurophysiology 27(1), 267–278. doi: 10.1152/jn.00108.2021.

Smith JC, Ellenberger HH, Ballanyi K, Richter DW & Feldman JL (1991). Pre-Botzinger complex: a brainstem region that may generate respiratory rhythm in mammals. Science 254, 726–729.

Smith JC, Abdala AP, Koizumi H, Rybak IA, Paton JF (2007). Spatial and functional architecture of the mammalian brain stem respiratory network: a hierarchy of three oscillatory mechanisms. Journal of Neurophysiology 98, 3370–87. doi: 10.1152/jn.00985.2007.

Smith JC, Abdala AP, Borgmann A, Rybak IA, Paton JF (2013). Brainstem respiratory networks: building blocks and microcircuits. Trends Neurosci. 36, 152–62. doi: 10.1016/j.tins.2012.11.004.

Takatoh J, Park JH, Lu J, Li S, Thompson P, Han B-X, Zhao S, Kleinfeld D, Friedman B & Wang F (2021). Constructing an adult orofacial premotor atlas in Allen mouse CCF ed. Chesler AT, Dulac C & Chesler AT. eLife 10, e67291.

Tan W, Janczewski WA, Yang P, Shao XM, Callaway EM, Feldman JL (2008). Silencing preBötzinger complex somatostatin-expressing neurons induces persistent apnea in awake rat. Nature Neuroscience 11, 538–40. doi: 10.1038/nn.2104.

Thoby-Brisson M, Ramirez JM (2001) Identification of two types of inspiratory pacemaker neurons in the isolated respiratory neural network of mice. Journal of Neurophysiology 86, 104–12.

Trevizan-Baú P, Dhingra RR, Furuya WI, Stanić D, Mazzone SB & Dutschmann M (2021). Forebrain projection neurons target functionally diverse respiratory control areas in the midbrain, pons, and medulla oblongata. Journal of Comparative Neurology 529, 2243–2264.

Yang CF, Kim EJ, Callaway EM & Feldman JL (2020). Monosynaptic Projections to Excitatory and Inhibitory preBötzinger Complex Neurons. Frontiers in Neuroanatomy 14, 58. doi: 10.3389/fnana.2020.00058.

